# Maize root-associated microbes likely under adaptive selection by the host to enhance phenotypic performance

**DOI:** 10.1101/2021.11.01.466815

**Authors:** Michael A. Meier, Gen Xu, Martha G. Lopez-Guerrero, Guangyong Li, Christine Smith, Brandi Sigmon, Joshua R. Herr, James R. Alfano, Yufeng Ge, James C. Schnable, Jinliang Yang

## Abstract

The root-associated microbiome (rhizobiome) plays a non-negligible role in determining plant health, stress tolerance, and nutrient use efficiency. However, it remains unclear to what extent the composition of the rhizobiome is governed by intraspecific variation in host plant genetics in the field and the degree to which host plant selection can reshape the composition of the rhizobiome. Here we quantify the rhizosphere microbial communities associated with a replicated diversity panel of 230 maize (Zea *mays* L.) genotypes grown in agronomically relevant conditions under high N (+N) and low N (-N) treatments. We show that the abundance of many root-associated microbes within a functional core microbial community of 150 abundant and consistently reproducible microbial groups is explainable by natural genetic variation in the host plant, with a greater proportion of microbial variance attributable to plant genetic variation in low N conditions. Population genetic approaches identify signatures of purifying selection in the maize genome associated with the abundance of several groups of microbes in the maize rhizobiome. Genome-wide association studies conducted using rhizobiome phenotypes identified n = 467 microbe-associated plant loci (MAPLs) in the maize genome linked to variation in the abundance of n = 115 microbial groups in the maize rhizosphere. In 62/115 cases, which is more than expected by chance, the abundance of these same microbial groups was correlated with variation in plant vigor indicators derived from high throughput phenotyping of the same field experiment. This study provides insights into harnessing the full potential of root-associated microbial symbionts in maize production.

## Introduction

Symbiotic relationships between plant hosts and root-associated microbes have been shaped through natural selection over millions of years of coevolution (Limborg and Heeb, 2018), and have been a driver of crop performance and yield in agricultural production since the beginning of plant domestication (Yadav et al., 2018). Microbial actors in the rhizosphere have been shown to promote plant growth (Saleem et al., 2019), improve nutrient use efficiency (Gomes et al., 2018; Zhu et al., 2016), and reduce abiotic stress response (Hussain et al., 2018). The promise of high throughput screens for plant growth promoting activity in isolated microbial strains or synthetic communities (Singer et al., 2021; Yee et al., 2021) is the potential discovery of microbial agents that can be used as seed or soil additives to improve crop performance under field conditions. Promising results have been observed in controlled environments (Van Gerrewey et al., 2020; Xi et al., 2020; Yu et al., 2021), but it remains a challenge to achieve similar outcomes in crops under agriculturally relevant field conditions (Eida et al., 2017; Kaur et al., 2020; Sessitsch et al., 2019). Many microbial inoculants struggle to compete with naturally occurring microbes in the rhizosphere and rarely survive for extended periods of time in the field (Piromyou et al., 2011). An improved understanding of how plants shape the composition of their rhizobiomes under diverse field conditions would make it more feasible to identify beneficial plant-microbe interactions that will be persistent and replicable in field environments. Moreover, studying the effects of plant genetics on microbial communities may identify opportunities to breed crop plants that recruit and maintain superior growth-conducive microbial communities from the natural environment.

Few studies to date have addressed the relationship between plant genetics and rhizobiomes in field settings, mainly because large-scale rhizosphere sampling (as opposed to leaf microbiome sampling) and DNA sequence analysis of microbial communities in diverse plant hosts is time-consuming, expensive, and poses significant logistical and technical challenges. It has been shown that plant genetics can explain variation in both root architecture (Bray and Topp, 2018) and exudation (Mönchgesang et al., 2016). If these factors in turn shape microbial communities (Sasse et al., 2018), variation in the plant-associated microbiota (hereafter referred to as rhizobiome traits) may also result from genetic factors. Recent studies suggested that the variation in the composition of root-associated microbiomes is likely controlled by plant genetic factors (i.e., heritable) in maize (Peiffer et al., 2013), sorghum (Deng et al., 2021), and switchgrass (Sutherland et al., 2021). However, to what extent these heritable microbes are affected by the plant host and contribute to variation in the crop phenotype remains unclear. Like any other trait under heritable genetic control, rhizobiome traits can be targeted in selective breeding experiments. To explore this idea, previous efforts have been directed towards identifying plant genetic loci that are associated with rhizobiome traits. Initial studies have shown that rhizosphere microbial communities differ between distinct genotypes of the same host species, which has been shown in a study on 27 maize genotypes (Peiffer et al., 2013; Walters et al., 2018) and more recently, in a diversity panel of 200 sorghum lines (Deng et al., 2021). Genome-wide association study (GWAS) has successfully revealed associations between plant genes and rhizobiome traits at high-level measures of rhizosphere community dissimilarity (i.e., using principal components) in an *Arabidopsis* diversity panel (Bergelson et al., 2019) or at order level (derived from operational taxonomic units (OTUs)) in a sorghum diversity panel (Deng et al., 2021). However, previous attempts at linking plant genes to the abundance of specific groups of microbes have had limited success due to small population size, limited host genetic diversity, or due to insufficient taxonomic resolution (Beilsmith et al., 2019; Liu et al., 2021). It was observed in previous studies (Meier et al., 2021; Zhu et al., 2016) that maize rhizosphere microbial communities drastically change in response to N fertilization or lack thereof. A possible explanation for this could be that the vast majority of the interval between maize domestication and the present, beneficial plant-microbe interactions have evolved in low-input agricultural systems characterized by relative scarcity of nutrients, predominantly nitrogen (N) (Brisson et al., 2019). This is in stark contrast to the modern agricultural environment that has been the norm since the 1960s, in which plants are supplied with large quantities of inorganic N fertilizer (Cao et al., 2018). As a consequence, previous selection pressure to retain traits favorable under low N conditions, including plant growth-promoting microbes, has been largely reduced in modern maize breeding programs (Haegele et al., 2013; Zhu et al., 2016). Thus, if a microbial group is indeed under host genetic control and has an effect on plant fitness (i.e., promotes plant development or increases crop yield) under either N condition, we would expect the rhizobiome trait to be under host selection.

In the present study, we evaluate the role that selection on plant genetic factors has played in shaping the maize root-associated microbiome under both current agricultural practices and under the historical low N regime. We employ the Buckler-Goodman diversity panel, a set of maize lines selected for maximum representation of genetic diversity and growth in temperate latitudes (Flint-Garcia et al., 2005). This population has previously been used to determine the heritability of leaf microbiome traits and to perform genome-wide association studies (GWAS) on a number of different phenotypes (Wallace et al., 2018). We collected replicated data on the root-associated microbiome of 230 lines drawn from this panel when grown under either high N (+N) and low N (−N) conditions in the field. For 150 rhizobiome traits, which were abundant and consistently reproducible, we quantify the degree to which variation is subject to plant genetic control, and test for evidence of selection under either or both N treatments. Using a set of 20 million high density single nucleotide polymorphisms (SNPs), we perform GWAS for each rhizobiome trait identifying genomic loci that are associated with one or more rhizobiome traits. Through comparison with gene expression data generated for the same population, we determine whether genes near microbe-associated plant loci are preferentially expressed in root tissue. Lastly, we evaluate whether the abundance of each microbial group in the rhizosphere is correlated with plant performance traits measured in the field, and whether microbe abundance and plant performance depend on the allele variant at selected microbe-associated plant loci. The results presented in this study lay the groundwork for future endeavors to investigate the molecular mechanisms of specific plant-microbe interactions under agronomically relevant conditions.

## Materials and Methods

### Field and experimental design

In this study, 230 maize (*Zea mays* subsp. *mays*) lines from the Buckler-Goodman maize association panel (Flint-Garcia et al., 2005) were planted in May of 2018 and 2019 in a rain-fed experimental field site at the University of Nebraska Lincoln’s Havelock Farm, at the geographic location [N 40.853, W 96.611]. In both years, the experiment followed commercial maize. Individual entries consisted of 2 row, 5.3 m long plots with 0.75 m alleyways between sequential plots, 75 cm spacing between rows, and 15 cm spacing between sequential plants. In each year, the experimental field was divided into 4 quadrants and the complete set of genotypes was planted in each quadrant following an incomplete block design (**Supplementary Figure 1**). N fertilizer (urea) was applied at the rate of 168 kg/ha to two diagonally opposed quadrants before planting, while two quadrants were left unfertilized (−N treatment).

### Rhizobiome sample preparation and sequencing

In 2018, n = 304 rhizosphere samples were collected from 28 maize genotypes (2 samples per subplot, 2 replicated plots per genotype and N treatment); and in 2019, n = 3,009 samples were collected from 230 genotypes (3 samples per subplot, 2 replicated plots per genotype and N treatment). Eight weeks after planting (2018: July 10 and 11; 2019: July 30, 31 and August 1), plant roots were dug up to a depth of 30 cm and rootstocks were manually shaken to remove loosely adherent soil. DNA was isolated using the MagAttract PowerSoil DNA KF Kit (Qiagen, Hilden, Germany) and the KingFisher Flex Purification System (Thermo Fisher, Waltham, MA, USA). DNA sequencing was performed using the Illumina MiSeq platform at the University of Minnesota Genomics Center (Minneapolis, MN, USA). In brief, 2×350 bp stretches of 16S rDNA spanning the V4 region were amplified using V4_515F_Nextera and V4_806R_Nextera primers, and the sequencing library was prepared as described by Gohl (Gohl et al., 2016).

### Raw read processing and construction of microbiome dataset

Paired-end 16S sequencing reads from a total of 3,313 samples were processed in R 3.5.2 using the workflow described by Callahan (Callahan et al., 2016a), which employs the package dada2 1.10.1(Callahan et al., 2016b). Taxonomy was assigned to amplicon sequence variants (ASVs) using the SILVA database version 138 (Yilmaz et al., 2014) as the reference. Raw ASV reads were subjected to a series of filters to produce a final ASV table with biologically relevant and reproducible 16S sequences (**Supplementary Dataset 1**). Only ASVs that were highly abundant and repeatedly observed in both years of sampling were considered for downstream analysis. ASVs were clustered into 150 functional groups of rhizosphere microbes using a procedure described previously (Meier et al., 2021). In short, each genus present in the dataset was analyzed separately and ASVs were clustered into groups that show consistent responses based on the number of total observations and based on the differential abundance under +N vs −N treatment (fold change of ASV counts), calculated using DESeq2 (Love et al., 2014).

### Heritability estimation

Heritability (h^2^) of microbial traits was calculated separately for +N and −N conditions using maize genotypes in the 2019 dataset for which balanced data was available. For each of the 150 rhizobiome traits, combined ASV counts were normalized by converting to relative abundance and subsequent natural log transformation. Using these transformed values, h^2^ was estimated following Deng et al. (Deng et al., 2021) for each microbial trait using R package sommer 4.1.0 (Covarrubias-Pazaran, 2016). In short, h^2^ is the amount of variance explained by the genotype term (V_genotype_) divided by the variance of the genotype and the error (V_genotype_ + V_error_/n), where n = 6 is the total number of samples (i.e., 2 replicates x 3 samples per replicate) used in this dataset. Heritability was tested for significance using a permutation test. For each trait the genotype labels of microbial abundance data were shuffled 1,000 times, and the distribution of heritabilities calculated from these shuffled datasets were used to assess the likelihood of observing the heritabilities calculated from the correctly labeled data under a null hypothesis of no host genetic control.

### Estimation of genetic architecture parameters

For the 150 rhizobiome traits, a Bayesian-based method (Zeng et al., 2018) was used to estimate genetic architecture parameters simultaneously, including polygenicity (i.e., proportion of SNPs with non-zero effects), SNP effects, and the relationship between SNP effect size and minor allele frequency. For the analysis, genotypic data of the maize diversity panel was obtained from the Panzea database and uplifted to the B73_refgen_v4 (Bukowski et al., 2018; Woodhouse et al., 2021). To account for SNP linkage disequilibrium (LD), a set of 834,975 independent SNPs (MAF >= 0.01) were retained by pruning SNPs in LD (window size 100 kb, step size 100 SNPs, *r*^*2*^ ≥ 0.1) using the PLINK1.9 software (Chang et al., 2015). In the analysis, the “BayesS” method was used with a chain length of 410,000 and the first 10,000 iterations as burnin.

### Genome-wide association study

We chose to use the best linear unbiased prediction (BLUP) of the natural log transformed relative abundance of ASV counts as the dependent variable for the GWAS analysis. Since only a fraction of genotypes were sampled from the 2018 field experiment, only sample data collected in 2019 was used for the BLUP calculation. A BLUP value was calculated for each microbial group and each treatment using R package lme4 (Bates et al., 2015). In the analysis, the following model was fitted to the data: Y ~ (1|genotype) + (1|block) + (1|split plot) + (1|split plot block) + error, where Y represents a rhizobiome trait (ln(ASV count of a microbial group / total ASV count in sample)) (see **Supplementary Figure 1** for more details of the field experimental design). GWAS was performed separately for each rhizobiome trait and for both the +N and −N treatment using GEMMA 0.98 (Zhou and Stephens, 2012) with a set of 21,714,057 SNPs (MAF >= 0.05) (Bukowski et al., 2018). In the GWAS model, the first three principal components and the kinship matrices were fitted to control for the population structure and genetic relatedness, respectively.

### RNA sequence analysis

Gene expression was analyzed using two independent datasets. The first dataset was obtained from Kremling (Kremling et al., 2018) and included RNA sequencing data from 7 different maize tissue types. The second RNA sequencing dataset was generated from root and leaf tissue collected 14 days after germination from both +N and −N treated pots using 4 genotypes from the maize diversity panel. Libraries were sequenced using the Illumina Novaseq 6000 platform with 150 bp paired-end reads. Sequencing reads were mapped to the B73 reference genome (AGP V4) (Jiao et al., 2017; Schnable et al., 2009) and gene expression was quantified using Rsubread (Liao et al., 2019).

### Phenotyping of plant traits

A total of 17 plant traits were measured in the 2019 field experiment. First, 15 leaf physiological traits were measured on the same days the rhizobiome samples were collected, and included leaf area (LA), chlorophyll content (CHL), dry weight (DW), fresh weight (FW), as well as concentrations of the elements B, Ca, Cu, Fe, K, Mg, Mn, N, P, S, and Zn. Measurement of the leaf traits was carried out as previously described (Ge et al., 2019). Two aerial imaging traits, canopy coverage (CC) and excess green index (ExG), were collected on August 12, 2019, 11-13 days after rhizobiome sample collection (Rodene et al., 2021).

### Data and code availability

The sequencing data reported in this publication (3,313 samples) can be accessed via the following five Sequence Read Archive (SRA) accession numbers: PRJNA771710, PRJNA771712, PRJNA771711, PRJNA685208, PRJNA685228 (summarized under the umbrella BioProject PRJNA772177). Scripts used to analyze the data are available on GitHub (https://github.com/mandmeier/HAVELOCK_BG).

## Results

### Characterization of the rhizobiome for diverse maize genotypes under two different N conditions

Paired-end 16S sequencing of 3,313 rhizosphere samples from 230 replicated genotypes of the maize diversity panel (Flint-Garcia et al., 2005) collected from field experiments conducted under both +N and −N conditions (**Materials and Methods, Supplementary Figure 1**) produced 216,681,749 raw sequence reads representing 496,738 unique amplicon sequence variants (ASVs) (**Materials and Methods**). Raw reads were subjected to a series of quality checks and abundance filters (**Supplementary Methods**) following a workflow for 16S sequencing data analysis by (Callahan et al., 2016a), which resulted in a curated dataset of 3,626 ASVs, 3,313 samples, and 105,722,181 total ASV counts (**Supplementary Dataset 1**). This dataset includes ASVs that are highly abundant across the maize diversity panel and reproducible in both years of sampling. Constrained Principal Coordinates analysis calculated from the abundances of 3,626 ASVs shows divergence of samples collected under either −N or +N treatment (**Figure 1A**), which indicates that the microbiomes differ between these two experimental conditions. The final 3,626 ASVs were clustered into n = 150 functionally distinct microbial groups (referred to as “rhizobiome traits”), spanning 19 major classes of rhizosphere microbiota (**Figure 1B, Supplementary Datasets 2 & 3**) using a method previously described (Meier et al., 2021). Of these rhizobiome traits, 79/150 (52.7%) groups were significantly more abundant in samples collected from the +N condition (t-test, p < 0.05), 53/150 (35.3%) significantly more abundant in samples collected from the −N condition, and 18/150 (12.0%) showed no significant difference in abundance between the two treatments. In several cases, more closely related microbial groups exhibit shared patterns of differential abundance between N treatments (**Supplementary Figure 2**).

**Figure 1:**
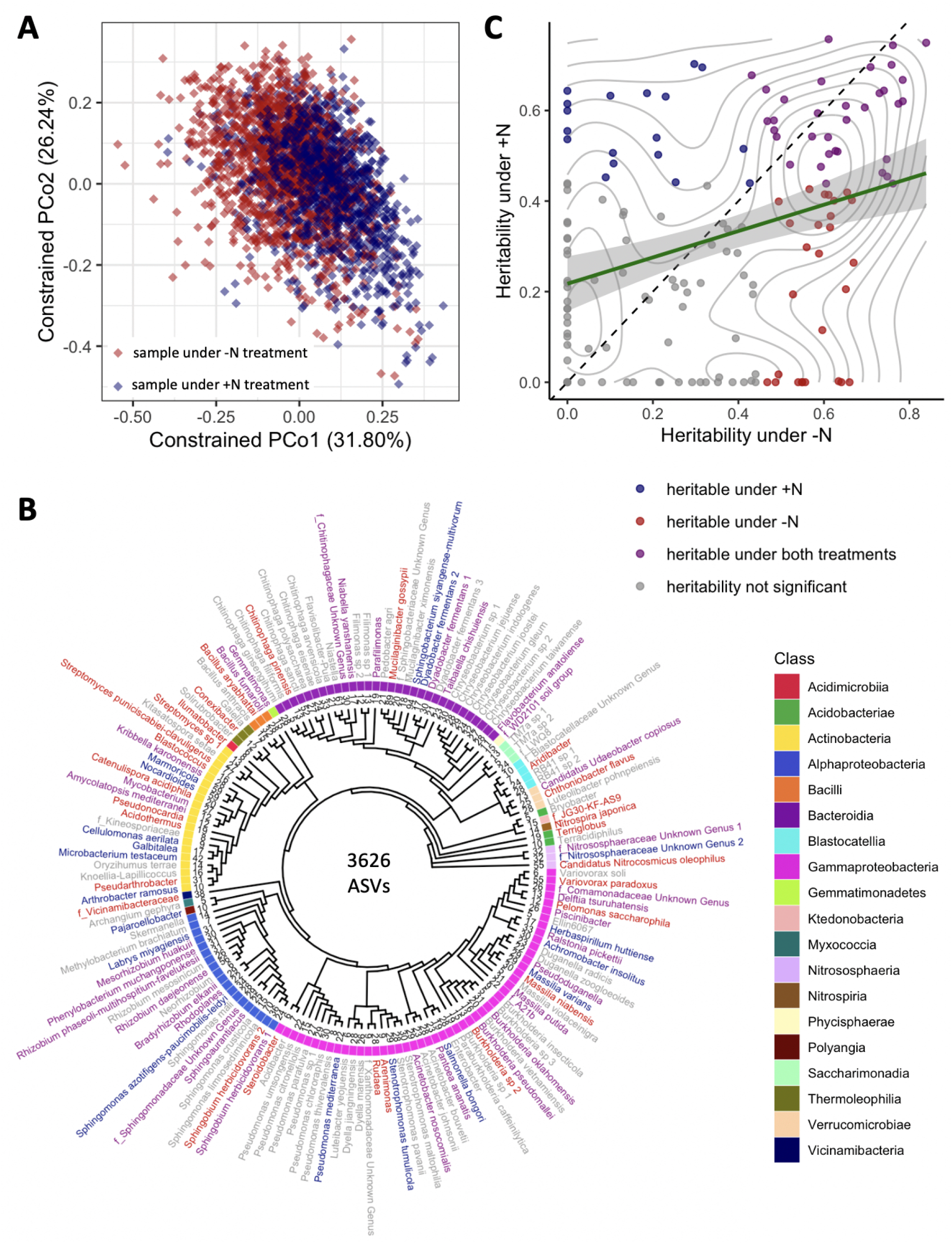
Diversity, phylogenetics, and heritability of rhizobiome traits. (**A**) Constrained ordination analysis showing the largest two principal coordinates calculated from the abundances of 3,626 ASVs. Each diamond represents one sample collected from plants under +N (blue) and −N (red) treatment, respectively. Note the separation of N treatments along PCo1. (**B**) Phylogenetic tree of 150 taxonomic groups of rhizosphere microbiota ( “rhizobiome traits”) generated by clustering 3,626 ASVs. Families are prefixed with “f_”, genus and species names are given where known. Numbers at tree tips indicate distinct ASVs in each group. Label colors indicate heritability of each rhizobiome trait as in panel C. (**C)** Heritability (h^2^) calculated for all 150 rhizobiome traits under +N and −N treatments. Green line indicates linear regression with 95% confidence interval, *r*^2^ = 0.104. Diagonal dashed line marks identity. Grey lines mark density of data points. Colors indicate whether traits are significantly heritable under either or both N treatments, as determined through a permutation analysis using 1000 permutations.

### Rhizobiome traits are more heritable under −N conditions

The abundance of each of the 150 rhizobiome traits was assessed separately for +N and −N conditions, and the heritability (proportion of total variance explicable by genetic factors) was estimated using an approach following Deng (Deng et al., 2021) (**Materials and Methods)**. Rhizobiome traits were comparatively more heritable under −N than +N conditions (**Figure 1C**). We found 34/150 (22.7%) microbial groups to be significantly heritable (permutation test, p < 0.05, **Materials and Methods**) under both N conditions, 18/150 (12%) only under +N conditions, and 27/150 (18%) only under −N conditions. Twelve rhizobiome traits exhibited estimated h^2^ > 0.6 in both +N and −N conditions (**Supplementary Figure 3**). These include 4 groups of ASVs assigned to the order *Burkholderiales* (the genus *Pseudoduganella*, an unknown genus in the *Comamonadaceae* family, the family *A21b*, and *Burkholderia oklahomensis*) and 2 groups in the *Sphingomonadales* order (*Sphingobium herbicidovorans 1* and an unknown genus in the *Sphingomonadaceae* family). Notably, closely related microbial groups did not exhibit obvious patterns of shared high or low estimated heritabilities (**Figure 1B**). As heritabilities and responses to treatments can vary considerably within families, genera, and lower taxonomic ranks, this underscores the importance of adequate taxonomic resolution when analyzing rhizosphere microbial communities.

### Rhizobiome traits are predominantly under purifying selection

Changes in the microbial community in response to N treatments (Meier et al., 2021; Zhu et al., 2016) **(Figure 1**) could be the result of altered selection pressure under the historic −N and the contemporary +N regimes, and such selection can happen either by purging deleterious alleles (purifying selection) or by elevating the frequencies of advantageous alleles (positive selection). To test this hypothesis, a Bayesian-based (Genome-wide Complex Trait Bayesian analysis, or GCTB) method was used to test for signatures of selection for each rhizobiome trait (**Materials and Methods**). A set of n = 834,975 independent SNPs was used to estimate their effects on 150 rhizobiome traits as well as 17 conventional plant traits collected from the same population in the same field experiments (**Materials and Methods, Supplementary Dataset 4**). Using the relationship between effects of non-zero SNPs and their minor allele frequencies (MAFs) as a proxy for the signature of selection (Zeng et al., 2018), the S parameter was jointly estimated from the GCTB analysis for both rhizobiome traits (**Figure 2A**) and plant traits (**Figure 2B**). According to Zeng (Zeng et al., 2018), if S = 0 (i.e., the posterior distribution of S is insignificantly different from zero), the SNP effect is independent of MAF, suggesting a neutral selection for the trait. If there is selection acting on the trait, the SNP effect can be positively (S > 0) or negatively (S < 0) related to MAF, indicating positive and purifying selection, respectively.

**Figure 2:**
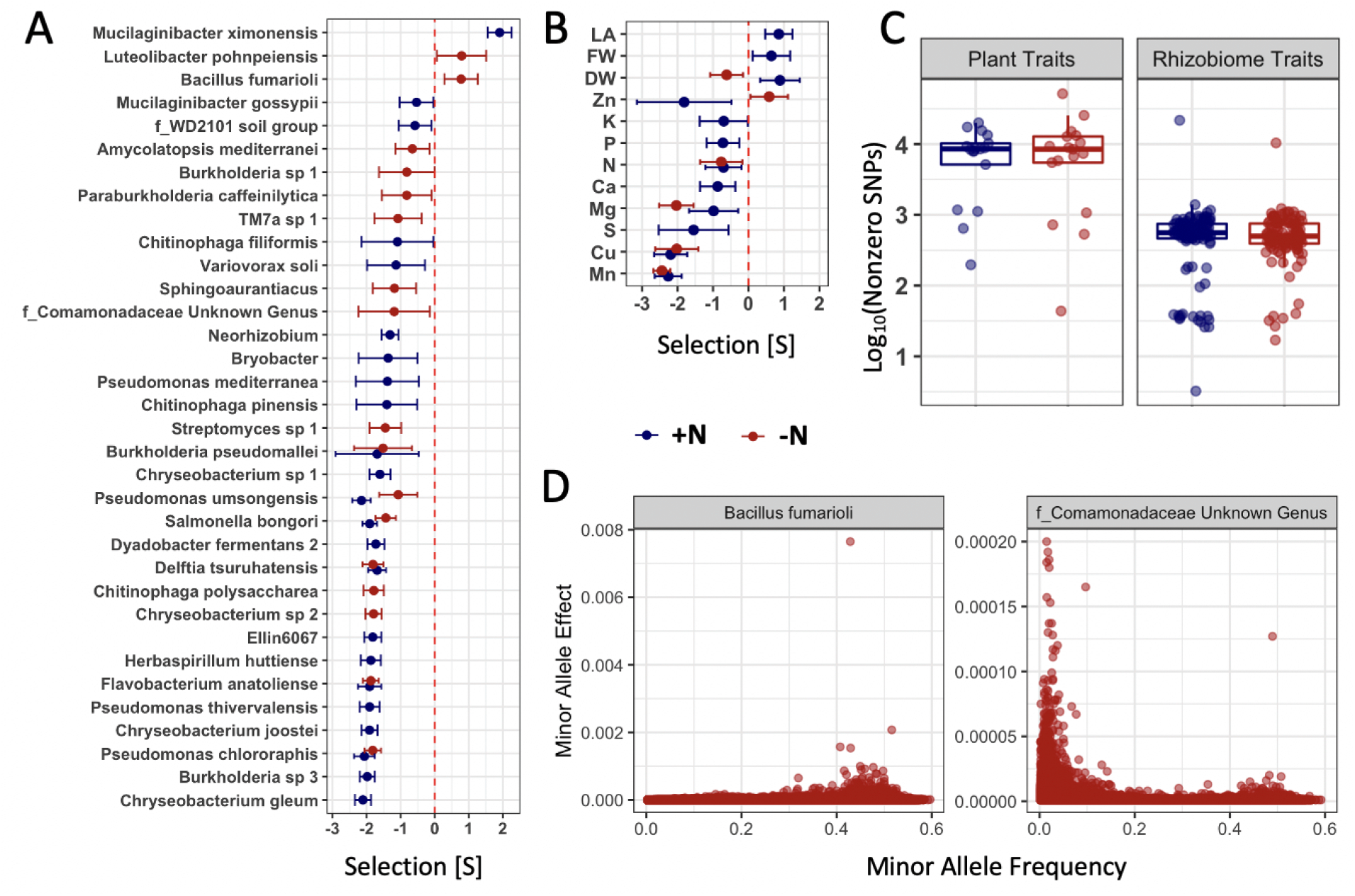
Population parameters estimated from genome-wide SNPs for plant and rhizobiome traits. Selection coefficients (S value) of rhizobiome (**A**) and plant (**B**) traits calculated for both N treatments using genome-wide independent SNPs. A negative S value indicates negative (purifying) selection, and a positive S value indicates positive (directional) selection. Traits are shown that show significant selection under one or both N treatments. (**C**) Number of SNPs showing non-zero effects for both plant and rhizobiome traits. (**D**) Examples of positive (*Bacillus fumarioli*) and purifying selection (*f_Comamonadaceae Unknown Genus*) showing minor allele effect vs. minor allele frequency with data skew to the right and to the left, respectively.

Several rhizobiome traits exhibited evidence of selection, and a substantial number of rhizobiome traits show evidence of purifying selection (22/150 under +N and 15/150 under −N). Three microbial groups (*Mucilaginibacter ximonensis*, *Luteolibacter pohnpeiensis*, *Bacillus fumarioli*) showed positive S values indicating that these traits might have been targets of positive selection. Relative to rhizobiome traits, plant leaf traits and nutrient traits were both more likely to exhibit evidence of selection within this maize population. Three out of 15 plant leaf traits, i.e., leaf area (LA), leaf fresh weight (FW), and leaf dry weight (DW) (**Materials and Methods**), exhibited S > 0 values under the +N condition, consistent with positive selection, while only one of the three exhibited a slightly negative S value in the −N condition and in that case exhibited a pattern consistent with weak purifying selection (**Figure 2B)**. Of the 11 micronutrient traits evaluated, 9/11 and 4/11 showed significantly negative S values in trait data collected under +N and −N conditions, respectively. From the same GCTB analysis, estimates of the number of SNPs with non-zero effects were substantially lower for rhizobiome traits than for conventional plant traits, whereas the differences were insignificant between the two N treatments for both rhizobiome and plant traits (**Figure 2C**). Using these non-zero effect SNPs, we plotted their minor allele frequency vs. the minor allele effect. As expected, in the case of positive selection (*Bacillus fumarioli*), we observed a skew towards higher MAF and in the case of purifying selection (*f_Comamonadaceae Unknown Genus*), a skew towards lower MAF (**Figure 2D)**.

### Genes underlying microbe-associated plant loci are preferentially expressed in root tissue

The observation that many rhizobiome traits are both under significant host genetic control and targets of selection suggests it may be possible to detect individual large effect loci controlling these rhizobiome traits. We performed GWAS using each of the 150 rhizobiome traits quantified under −N or +N conditions, to find specific associations between plant genomic loci and rhizobiome traits. We focused on “hotspots” along the genome where we find the highest cumulative density of significant associations between SNPs and any rhizobiome traits under either N treatment to mitigate false discoveries given the large number of total GWAS conducted. For this purpose, we tallied the number of significant (p < 10^−5^) GWAS signals in 10 kb genomic windows and defined the 75% quantile of all genomic windows containing at least one significant signal as microbe-associated plant loci (MAPLs) (**Materials and Methods**).

Using our arbitrary definition, a total of 467 MAPLs were detected across the maize genome, 246 under +N treatment and 221 under −N treatment (**Figure 3A, Supplementary Dataset 5**). Of the 150 rhizobiome traits evaluated here, 115 were associated with at least one MAPL in at least one of the two N treatments, with 74 rhizobiome traits associated with 221 MAPLs under the +N treatment and 90 rhizobiome traits with 245 MAPLs under the −N treatment. 49 rhizobiome traits showed significant associations under both N treatments, albeit with different plant loci. No loci were found that had associations with rhizobiome traits under both N treatments, which may suggest that plant gene - microbe interactions are dynamic and dependent on external factors or simply reflect limited statistical power to detect true associations given our limited number of genotypes and replicates.

**Figure 3:**
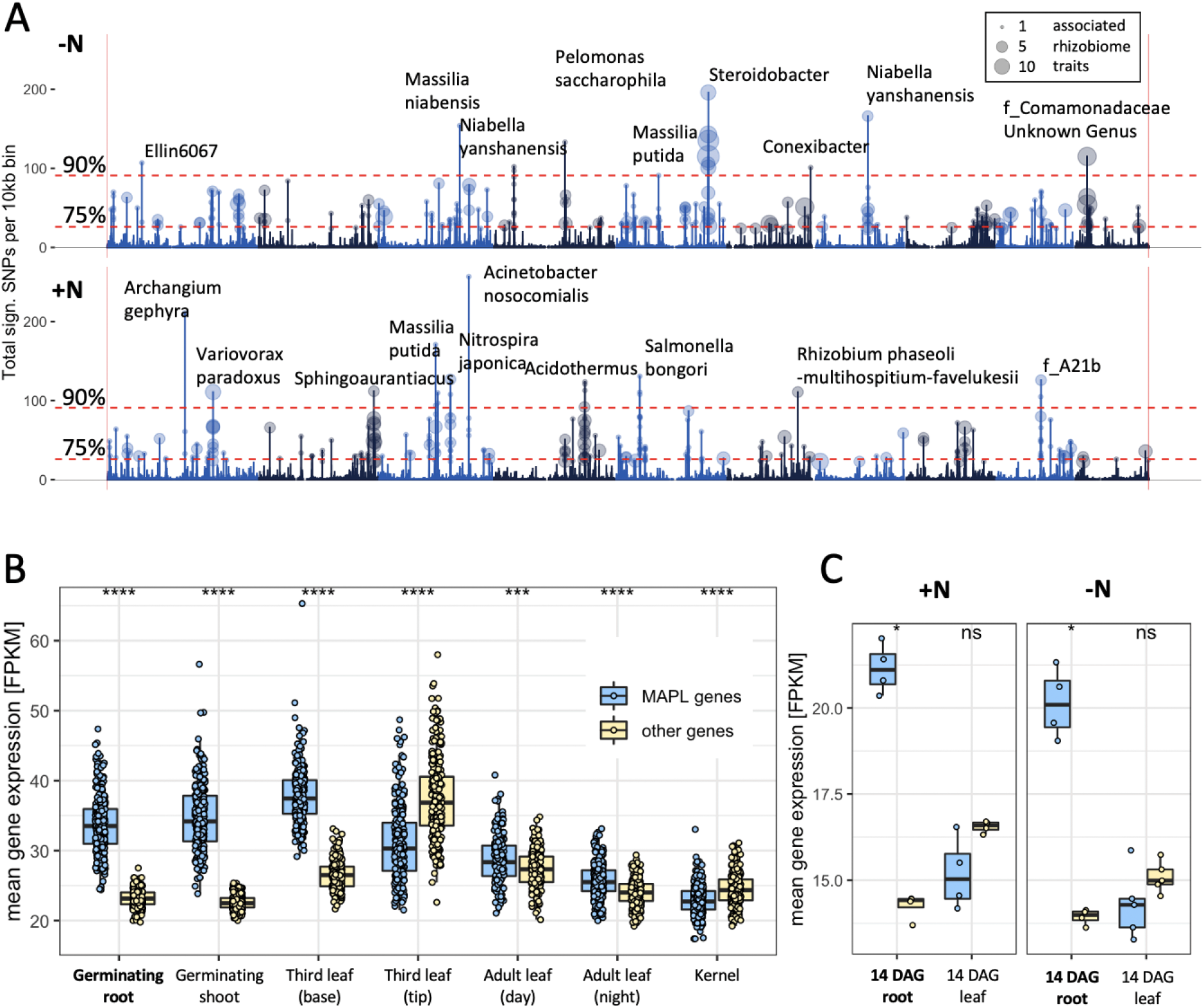
Microbe associated plant loci (MAPLs) contain genes expressed in roots. (**A)** GWAS of 150 rhizobiome traits reveals microbe-associated plant loci across the maize genome. Dashed lines indicate the 75% and 90% quantiles, respectively, to identify hotspots with particularly strong GWAS signals. Circles on top of peaks at each MAPL indicate the number of rhizobiome traits associated with each locus (traits that have at least one SNP above the 10^−5^ significance level). The dominant rhizobiome trait with the highest count of significant SNPs was annotated for each MAPL in the 90% quantile. (**B**) Mean gene expression of 305 MAPL genes and 29,540 other genes in seven tissue types, measured in 298 genotypes of the maize diversity panel (Kremling et al., 2018). (**C**) Mean gene expression of 395 MAPL genes and 43,751 other genes in two tissue types, measured in the present study in four maize genotypes under +N and −N treatments.

We hypothesized that many plant genes underlying MAPL hotspots may exert control over the rhizosphere microbiome via changes in root physiology, architecture, and exudate composition (Vandenkoornhuyse et al., 2015) and may therefore be preferentially expressed in root tissue. Transcribed sequences of 395 gene models were completely contained within ±10 kb of the 467 MAPL hotspots identified here, where 267/467 MAPLs contained between 1 and 6 genes. We evaluated the expression of these MAPL genes relative to the overall patterns exhibited by all genes outside the MAPL regions in seven tissues using published expression data from the same maize population (Kremling et al., 2018). Expression data was available in this dataset for 305 out of 395 MAPL genes across 298 maize genotypes. These MAPL genes, when compared to (n = 29,539) other genes available in the dataset, show on average significantly higher expression in the germinating root, the germinating shoot, and the third leaf base (**Figure 3B**).

We obtained additional RNA sequencing data from root and leaf tissue of two-week old seedlings derived from four maize genotypes (B73, SD40, K55, and W153R) under both N treatments. A similar pattern was observed in this dataset, with significantly higher expression of 395 MAPL genes in root but not leaf tissue compared to (n = 43,751) other genes available in this dataset (**Figure 3C**). This pattern was apparent under both N treatments, while no significant differences were observed in the pattern of MAPL gene expression between the two N treatments.

Collectively, these data are consistent with the hypothesis that root-associated microbial communities are at least in part genetically controlled by the host plant in a process mediated by root gene expression.

### Heritable and adaptively selected rhizobiota are associated with plant phenotypes

We investigated the correlation of microbe abundance with 17 plant traits, including leaf physiology, leaf micronutrient traits, and traits extracted from aerial images (**Materials and Methods**) to identify potential plant phenotypic consequences of variation in the abundance of specific rhizosphere microbes. Several rhizobiome traits were significantly correlated (p < 0.01) with measures of plant performance, such as leaf area, leaf dry weight and fresh weight, and with several measures of leaf micronutrients such as nitrogen, sulfur, and phosphorus (**Supplementary Figure 4**). The trait that was most strongly linked to microbe abundance was leaf canopy coverage (CC). In total, 62 microbial groups – more than expected by chance (permutation test p < 0.001) – were significantly (correlation p < 0.01) associated with CC, 15 under +N treatment, 18 under −N treatment, and 29 under both treatments (**Figure 4A**). 30 traits under +N and 35 under −N were positively correlated with CC. 14 traits under +N and 12 under −N were negatively correlated with CC. Among the same microbial groups, 44/62 (71%) showed significant evidence of host plant genetic control under either or both N treatments and 56/62 (90%) are associated with 255/395 (65%) genes in 174/467 MAPLs (39%) identified across the maize genome (**Supplementary Figure 5**). Under both N treatments, we observe an association between heritability and the correlation with CC, which was statistically significant (*r* = 0.39, p = 4×10^−6^) for +N and even more significant (*r* = 0.49, p = 1.7×10^−9^) under the −N regime (**Figure 4B**).

**Figure 4:**
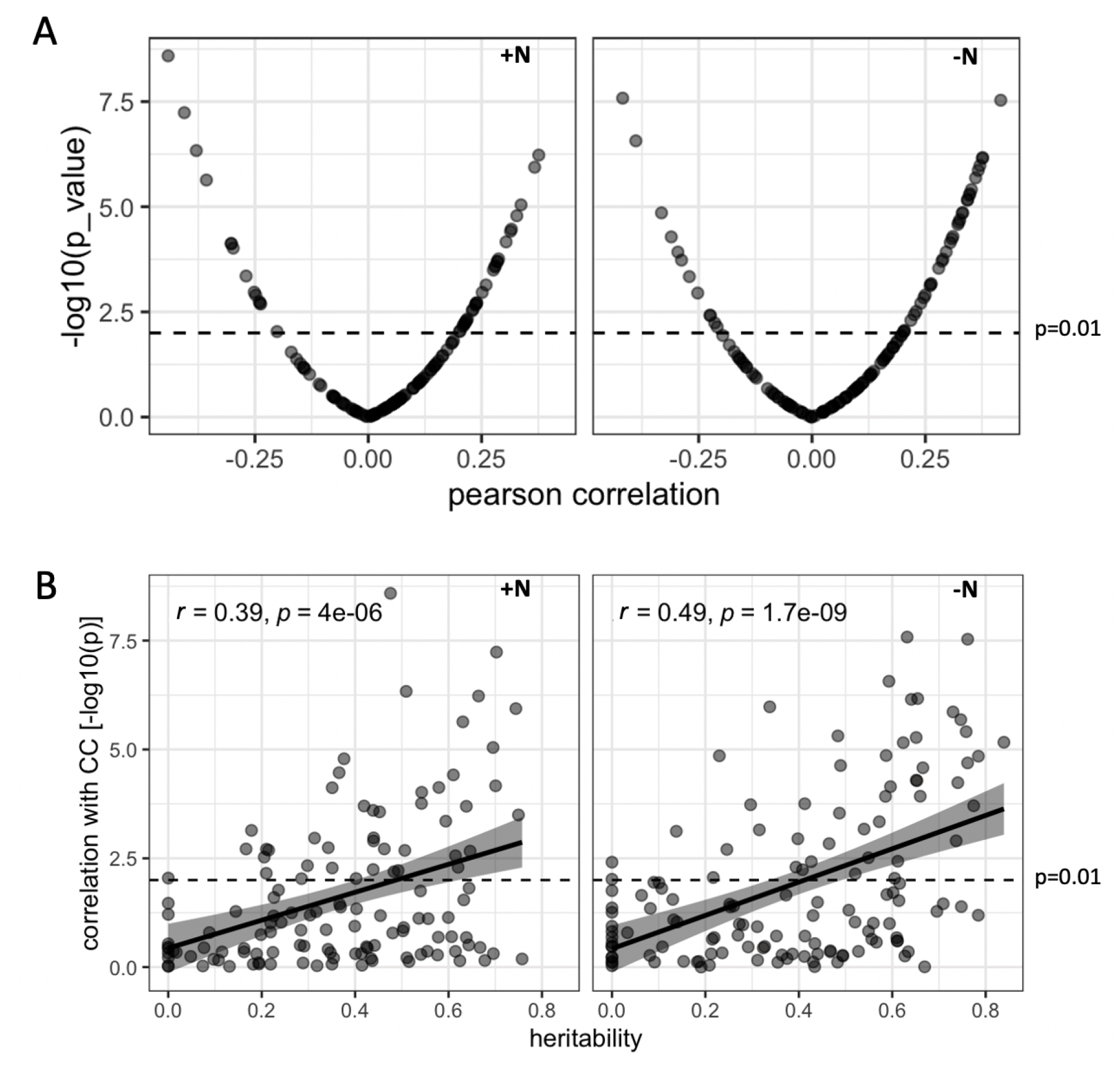
Heritable rhizobiome traits tend to be correlated with whole plant canopy coverage. (**A**) Distribution of statistical significance and correlation values for the relationship between canopy coverage (CC) and each of 150 rhizobiome traits under either +N or −N conditions. Dashed line indicates significance level (*p* = 0.01). (**B**) Relationship between the estimated heritability of individual rhizobiome traits and correlation of the same individual rhizobiome traits with variation in CC. Dashed line indicates significance level (*p* = 0.01).

### Allelic differences at microbe-associated plant loci predict microbe abundance

We identified several strong GWAS signals that link plant genomic loci to rhizobiome traits (**Figure 3A**). Such signals indicate that the pattern of SNPs at a given locus (i.e., the genetic architecture) has a large magnitude of effect attached to the abundance of the associated microbial groups. Next, we sought to determine whether a particular allele (either the major or the minor variant) in our maize population is associated with an increased or decreased abundance of the corresponding microbe.

The unknown genus in the *Comamonadaceae* family mentioned above, while unnamed and uncharacterized, shows high heritability under both N treatments (h^2^ = 0.610 under +N, and 0.651 under −N, **Figure 1B & 1C**), and shows evidence of being under purifying selection under −N (**Figure 2A & 2D**). Under the same environmental conditions, a significant MAPL controlling variation in microbial abundance is detectable on maize chromosome 10 (**Figure 3A** and **Figure 5A**). This same rhizobiome trait is among those that are positively correlated with CC under both −N (r = 0.347, p = 5.313×10^−6^) and +N (r = 0.314, p = 3.845×10^−5^) (**Figure 4A**). A total of five annotated gene models are located near the peak of significant SNP markers that define the chromosome 10 MAPL for this rhizobiome trait (**Figure 5A & 5B**). A linkage disequilibrium block was observed between 23.90 and 23.96 MB on maize chromosome 10, spanning the set of significantly associated SNPs and three candidate genes Zm00001d023838, Zm00001d023839 and Zm00001d023840 (**Figure 5C**). As described above, the abundance of the *f_Comamonadaceae* genus was significantly correlated with variation in CC, shown here for the −N treatment (**Figure 5D**). Next, we used the haplotype information at the target SNP to mark genotypes that carry the major allele or the minor allele, respectively, and the abundance of the *f_Comamonadaceae* genus was significantly higher in rhizosphere samples collected from maize genotypes carrying the major allele than in samples collected from maize genotypes carrying the minor allele (**Figure 5E)**. However, CC was not significantly different between maize genotypes carrying either the major or minor allele of the chromosome 10 MAPL (**Figure 5F**).

**Figure 5:**
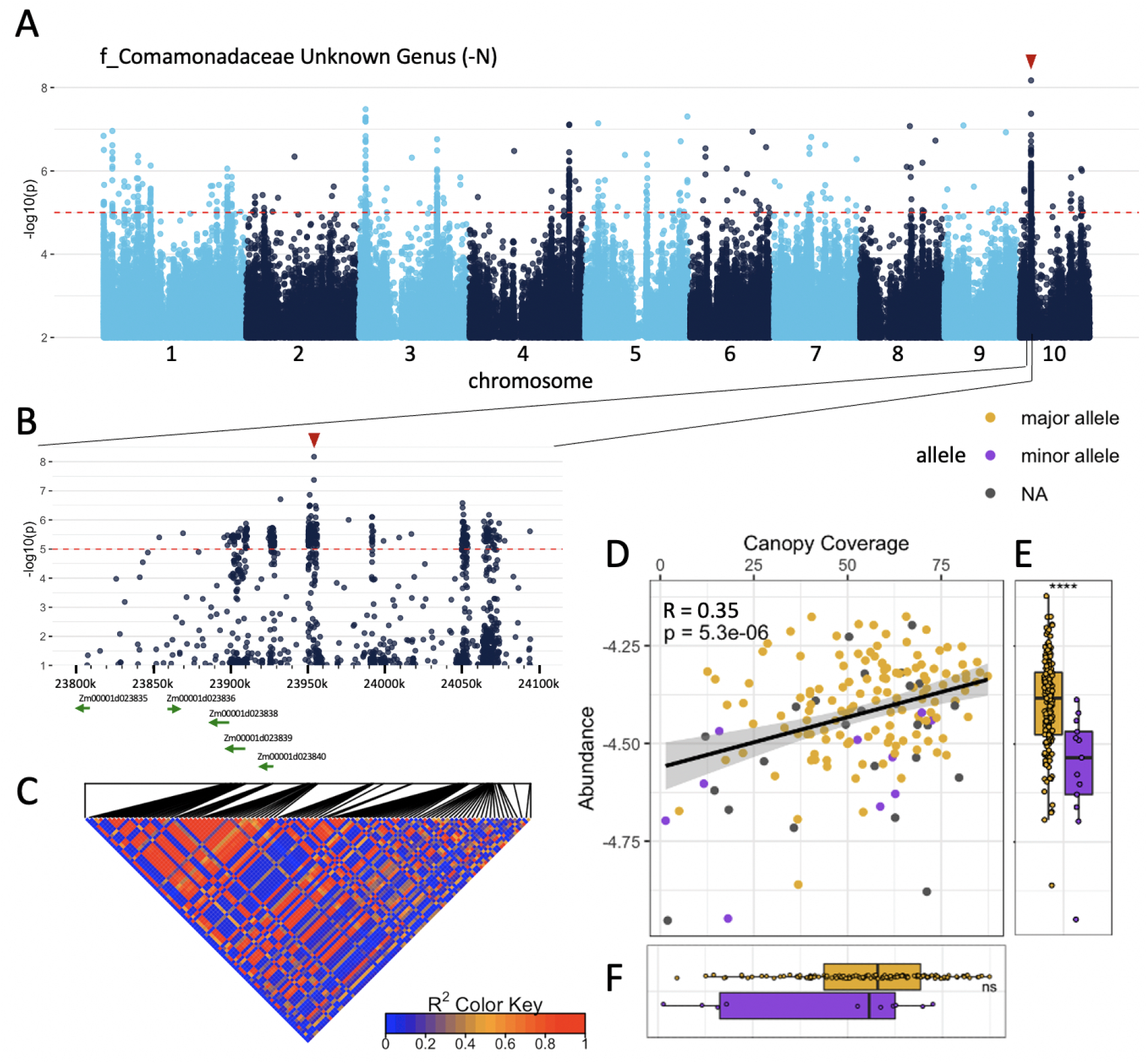
Abundance of heritable, adaptively selected microbes depends on allelic differences at MAPLs. (**A**) Results of a genome wide association study conducted using values for the rhizobiome trait (*f_Comamonadaceae Unknown Genus*) observed for ~230 maize lines grown under nitrogen deficient conditions. Alternating colors differentiate the 10 chromosomes of maize. Dashed line indicates a statistical significance cutoff of p = 10^−5^. (**B**) Zoomed in visualization of the region containing the peak observed on chromosome 10. (**C**) Linkage disequilibrium among SNP markers genotyped within this region, calculated using genotype data from 271 lines (**D**) Correlation plot of microbe abundance vs. canopy coverage (CC). Each point represents a maize genotype. Differences in microbe abundance (**E**) and CC (**F**) are marked between genotypes carrying the major allele (gold) vs the minor allele (purple) at the target SNP (red arrow in panel A and B).

## Discussion

This study profiled the rhizosphere inhabiting microbiota of several hundred maize genotypes under agronomically relevant field conditions. Through a 16S rDNA-sequencing based approach, we identified a set of 150 reproducible rhizobiome traits. In total, 79 out of the 150 rhizobiome traits (52%) showed significant evidence of being influenced by host plant genotype in one or more environmental conditions. The estimated heritability of rhizobiome traits in this study ranged from 0 to 0.757 for the +N treatment (mean 0.320) and from 0 to 0.839 for the −N treatment (mean 0.352). A comparable study of the root-associated microbiomes of the genotypes of a sorghum diversity panel estimated similar values (Deng et al., 2021). A previous study on the maize diversity panel (Wallace et al., 2018) investigated the heritability of 185 individual OTUs and 196 higher taxonomic units measured in the leaf microbiome. The study reported only 2 OTUs and 3 higher taxonomic groups showing significant heritability using the same permutation test we employed in this study. This may indicate that plant genetics have a stronger influence on rhizosphere colonizing microbes than on leaf colonizing microbes. One reason for this may be that there is a direct pathway for plant-to-microbe communication via root exudates (Doornbos et al., 2012). In contrast, no equivalent exchange of chemical information has been reported above ground, with the possible exception of aerial root mucilage (Van Deynze et al., 2018).

We observed relatively higher heritability for rhizobiome traits quantified from plants grown in the −N treatment than under the +N treatment. This outcome is consistent with a model where the partnerships between microbiomes and plants were established in natural and early agricultural systems which were predominantly N limited (Brisson et al., 2019). N insufficiency in maize induces dramatic changes in physiology (Ciampitti et al., 2013), gene expression (Chen et al., 2011), root architecture (Gaudin et al., 2011) and root exudation (Baudoin et al., 2003; Haase et al., 2007). Consistent with this, N fertilization or the lack thereof has substantial consequences on plant-microbe associations. In this study, 12% of rhizobiome traits were only significantly heritable under the +N treatment, and 18% only under the −N condition, and GWAS revealed plant-microbe associations at different genomic loci depending on the N treatment. Previous observations have also reported that rhizosphere microbial communities are highly sensitive to environmental conditions, in particular to N deficiency (Meier et al., 2021; Zhu et al., 2016). This finding emphasizes the need to optimize microbial communities not only for a specific host but also for specific levels of N fertilization.

Our results suggest that the capacity of maize plants to encourage or discourage colonization of the rhizosphere by specific microbiota has been a target of selection, in particular, we observed purifying selection acting on genetic variants associated with the abundance of 22 and 15 rhizosphere traits in the +N and −N environments, respectively. Notably, among the rhizosphere denizens whose abundance showed the strongest evidence of being a target of purifying selection in the host genome are 3 species of *Chitinophaga*, 4 species of *Chryseobacterium*, 4 species of *Pseudomonas* and 4 species of *Burkholderia*. Previous studies have reported plant growth promoting capability of various species of *Chitinophaga* (Sharma et al., 2020), *Chryseobacterium* (Dardanelli et al., 2010; Singh et al., 2013), *Pseudomonas* (Otieno et al., 2015; Preston, 2004) and *Burkholderia* (Bernabeu et al., 2015; Kurepin et al., 2015). Evidence of positive selection was observed for genetic variants in maize associated with variation in the abundance of *Bacillus fumarioli*, *Luteolibacter pohnpeiensis* and *Mucilaginibacter xinmonensis*. *B. fumarioli* has previously been observed in plant rhizospheres, particularly in maize (Garbeva et al., 2008), and several strains of *Bacillus* (Kumar et al., 2012) and *Mucilaginibacter* (Madhaiyan et al., 2010) have shown plant growth promoting activities. Selection of members of the *Luteolibacter* genus in plant rhizospheres has been directly observed in inoculation experiments of culturable isolates (Nunes da Rocha et al., 2013). Together, these findings indicate that the microbial groups we identified as under host plant genetic control may have a possible role in determining plant fitness.

Among the 150 rhizobiome traits analyzed here, 62 showed a significant correlation with measurements of canopy coverage collected from the same field experiment. In particular, the observed link between heritability of microbes and correlation with plant performance may indicate a symbiotic relationship of the host plant and root-associated microbes. Indeed, the majority of rhizobiome traits that are correlated with canopy coverage are both heritable and associated with one or more microbe-associated plant loci (MAPLs). Genes linked to variation in rhizobiome traits via GWAS were preferentially expressed in roots across genotypes in multiple independent gene expression datasets, suggesting a number of potential mechanisms for host plant genotypes to influence the composition of the rhizobiome. Altering the expression of particular MAPL genes identified here may be an avenue to directly influence and engineer the abundance of targeted microbial groups in the plant rhizosphere, although substantial further experimentation and study remains necessary.

We evaluated associations between rhizobiome traits and a number of whole plant phenotypes. However, the Buckler Goodman diversity panel has been and continues to be utilized in field experiments to determine the genetic basis of many phenotypes across diverse environments (Flint-Garcia et al., 2005; Ge et al., 2019; Kremling et al., 2018; Rice et al., 2020). The datasets generated here (**Supplementary Datasets**) link the abundance of 150 microbial groups in the rhizosphere to genetic variation in 230 genotypes across two N treatments. Combining these public datasets with plant phenotypes collected from the same genotypes in additional environments may lead to the identification of other cases where MAPLs are associated with variation in plant phenotypes or plant performance. The results presented in this study add to an increasing body of evidence (Bakker et al., 2013; Pérez-Jaramillo et al., 2016) that microbial communities are actively and dynamically shaped by host plant genetic variation and may serve as the foundation for future research into particular plant-microbe relationships that may be harnessed to sustainably increase crop productivity and resilience to abiotic stress.

## Acknowledgments

This study is supported by National Science Foundation EPSCoR Cooperative Agreement OIA-1557417. In Memory of James R. Alfano we thank him for his initiative and leadership at the Center for Root and Rhizobiome Innovation (CRRI). We also thank Edgar Cahoon and the CRRI team for setting up and maintaining an exemplary collaborative environment. We further acknowledge Tom Clemente, Karin van Dijk, Daniel Schachtman, Ellen Marsh, Lisa Vonfeldt, Alan Muthersbaugh, Jenny Stebbing and TJ McAndrew, for technical support. Lastly, we thank the many scientists who assisted in collecting rhizosphere samples: Laure-Olivia Mbouang Angoua, Bryce Askey, Abbas Atefi, Natalie Belz, Erin Bertone, Eledon Beyene, Alexandra Bradley, Amanda Butera, Christian Butera, Madelyn Calvert, Noah Carroll, Jessica Chen, Sierra Conway, Floreana Cordova, Xiuru Dai, Semra Palali Delen, Yavuz Delen, Tessa Durham-Brooks, Samuel Eastman, Alex Enerson, Ashley Foltz, Nick Friedman, Cierra Goerish, Wihib Hankore, Davron Hanley, Aris Hino, Chun Yin Ho, J. Preston Hurst, Kylie Irene, Panya Kim, Nataniel Korth, Courtney Krsnak, Enzo Lamontia, Zhikai Liang, Xiangdong Liu, Angelique Malcolm, Rajan Mediratta, Chenyong Miao, Xiaoxi Meng, Levi Nigro, Alejandro Pages, Connor Pedersen, Nathaniel Pester, Sam Polk, Raghuprakash Kastoori Ramamurthy, Eric Rodene, Daniel Santano de Carvalho, Emma Sheridan, Aris Shino, Isabel Sigmon, Taylor Stratman, Guangchao Sun, Michael Tross, Misty Wehling, Florian Wurtele and Zhikai Yang.

