## Supplementary Materials for "Maize root-associated microbes likely under adaptive selection by the host to enhance phenotypic performance"

#### Supplementary Datasets

- **Supplementary Dataset 1:** Feature table (3,313 samples by 3,626 ASVs) from which our results were generated, alongside the sample metadata collected in this study.
- **Supplementary Dataset 2:** Taxonomically annotated list of 3,626 16S sequences that comprise the core maize microbiome used for this analysis and may serve as a reference to identify the same maize-associated ASVs in future experiments. For description and units of plant traits refer to dataset 4.
- **Supplementary Dataset 3:** List of the 150 rhizobiome traits defined in this study alongside relevant summary statistics, such as abundance, heritability, selection coefficients, and correlations with plant traits under both N treatments.
- **Supplementary Dataset 4:** List of 229 Buckler-Goodman maize genotypes with the corresponding measurements of all 17 plant and 150 rhizobiome traits analyzed here under both N treatments. Sample-level data is published for aerial imaging (Rodene et al., 2021).
- **Supplementary Dataset 5:** List of 467 microbe-associated plant loci with associated rhizobiome traits and gene models.

### Supplementary Figures

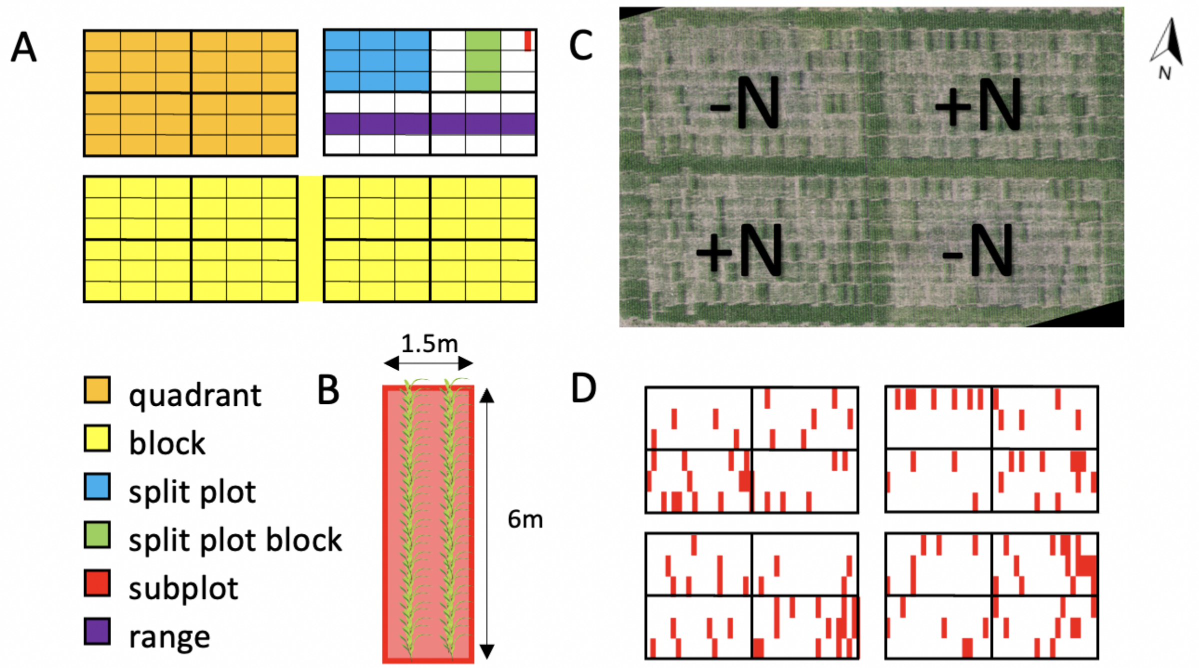


**Supplementary Figure 1: Field experimental design**. (A) Up to 230 maize genotypes were represented in each of 4 quadrants in 2 replicate blocks. Quadrants were planted in 6 ranges and divided into 4 split plots. Each split plot was divided into 3 split plot blocks, and each split plot block was divided into 21 subplots for a total of 252 subplots per quadrant. (B) Each 1.5m (5 ft) x 6m (20 ft) subplot (experimental unit) consisted of two rows of 36 maize plants of the same genotype, with a spacing of 75 cm (30 in) between rows and 15 cm (6 in) between plants. (C) Photomosaic of the 2019 field at flowering time. N fertilizer was applied to the NE and SW quadrants before planting. (D) 128 subplots across the field (marked in red) were planted with a check genotype (B73xMo17) in order to be able to quantify and control for spatial variation.

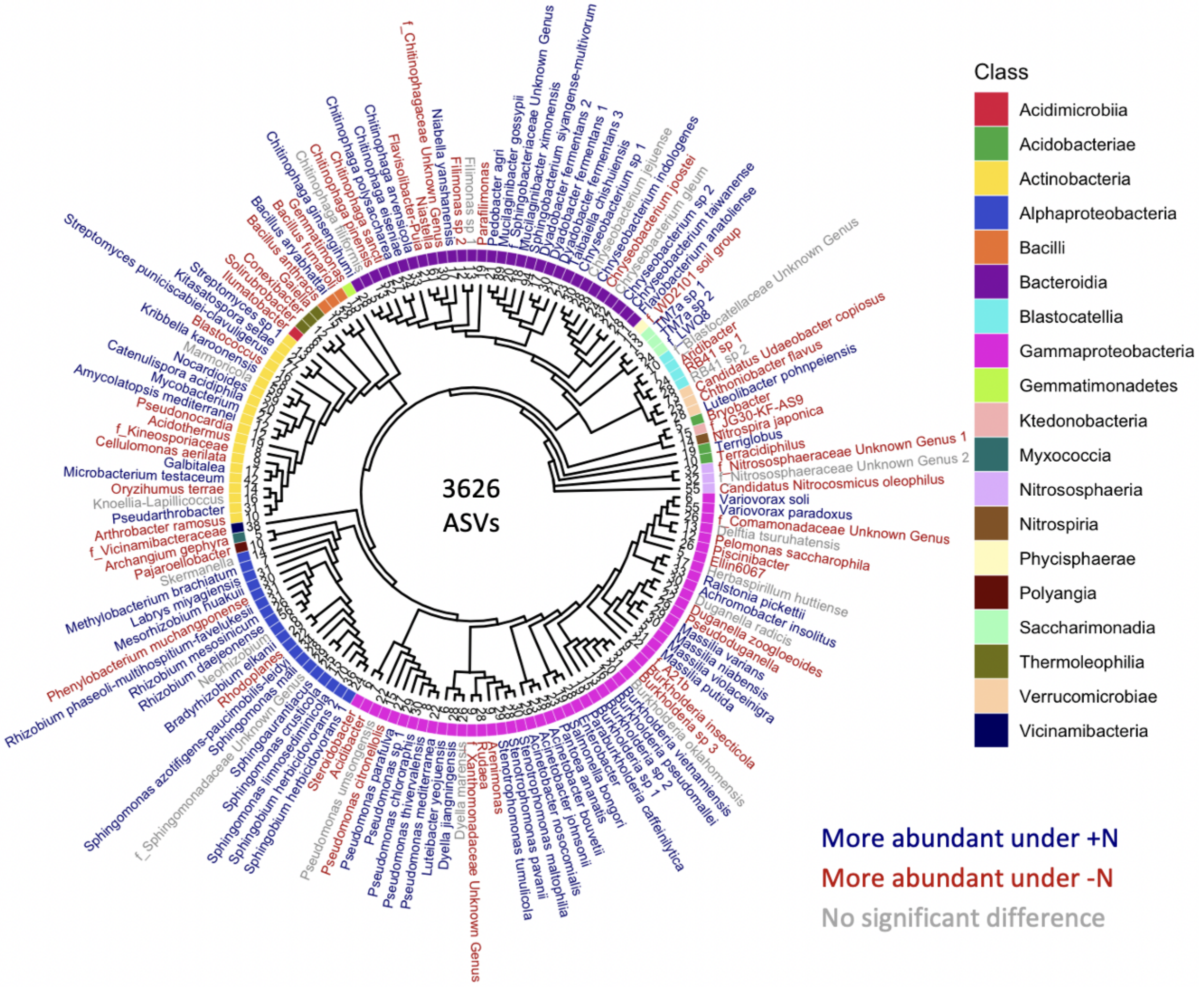


**Supplementary Figure 2**: **Phylogenetic tree of 150 microbial groups.** Colors indicate differential abundance between the +N and -N treatment.

**A**

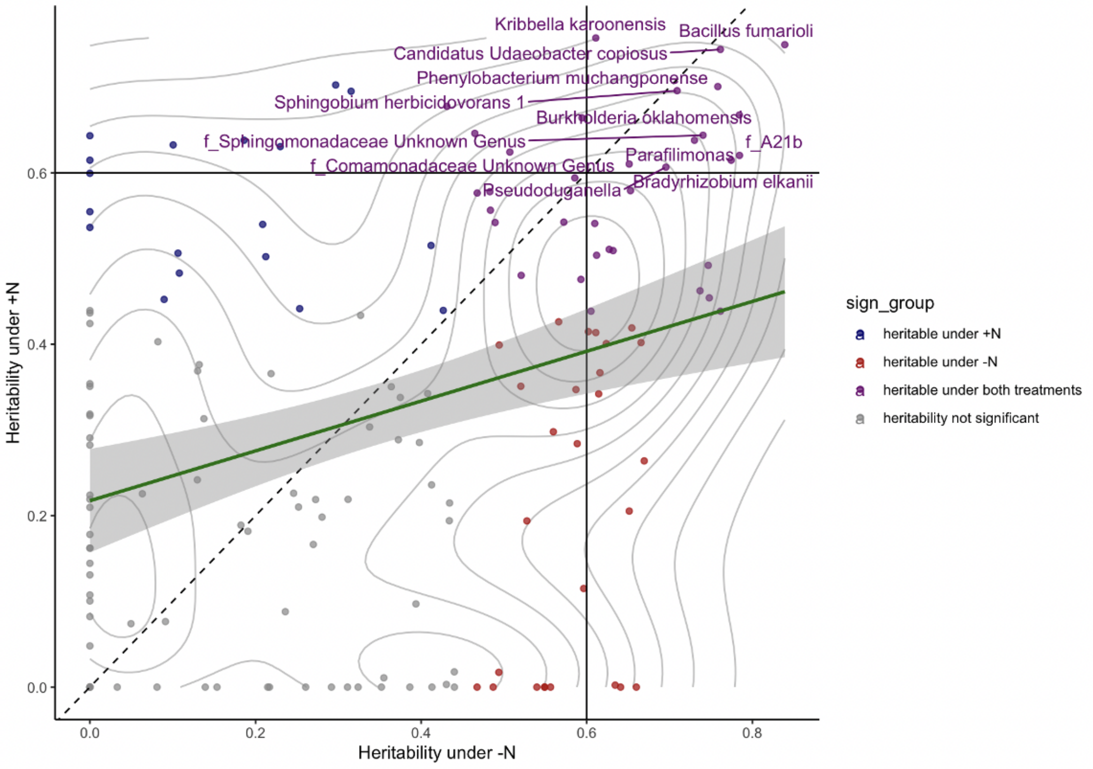


**B**

| Phylum | Class | Order | Family | tax_group |
|---|---|---|---|---|
| Verrucomicrobiota | Verrucomicrobiae | Chthoniobacterales | Chthoniobacteraceae | Candidatus Udaeobacter copiosus |
| Proteobacteria | Alphaproteobacteria | Sphingomonadales | Sphingomonadaceae | Sphingobium herbicidovorans 1 |
| Proteobacteria | Alphaproteobacteria | Sphingomonadales | Sphingomonadaceae | f_Sphingomonadaceae Unknown Genus |
| Proteobacteria | Alphaproteobacteria | Rhizobiales | Xanthobacteraceae | Bradyrhizobium elkanii |
| Proteobacteria | Alphaproteobacteria | Caulobacterales | Caulobacteraceae | Phenylobacterium muchangponense |
| Proteobacteria | Gammaproteobacteria | Burkholderiales | Oxalobacteraceae | Pseudoduganella |
| Proteobacteria | Gammaproteobacteria | Burkholderiales | Comamonadaceae | f_Comamonadaceae Unknown Genus |
| Proteobacteria | Gammaproteobacteria | Burkholderiales | A21b | f_A21b |
| Proteobacteria | Gammaproteobacteria | Burkholderiales | Burkholderiaceae | Burkholderia oklahomensis |
| Firmicutes | Bacilli | Bacillales | Bacillaceae | Bacillus fumarioli |
| Bacteroidota | Bacteroidia | Chitinophagales | Chitinophagaceae | Parafilimonas |
| Actinobacteriota | Actinobacteria | Propionibacteriales | Nocardioidaceae | Kribbella karoonensis |

**Supplementary Figure 3**: **Annotations of heritable microbial groups.** (A) The 12 most heritable microbial groups with heritability > 0.6 (drawn lines) under both N conditions were annotated by name. (B) Taxonomy of the 12 most heritable groups.

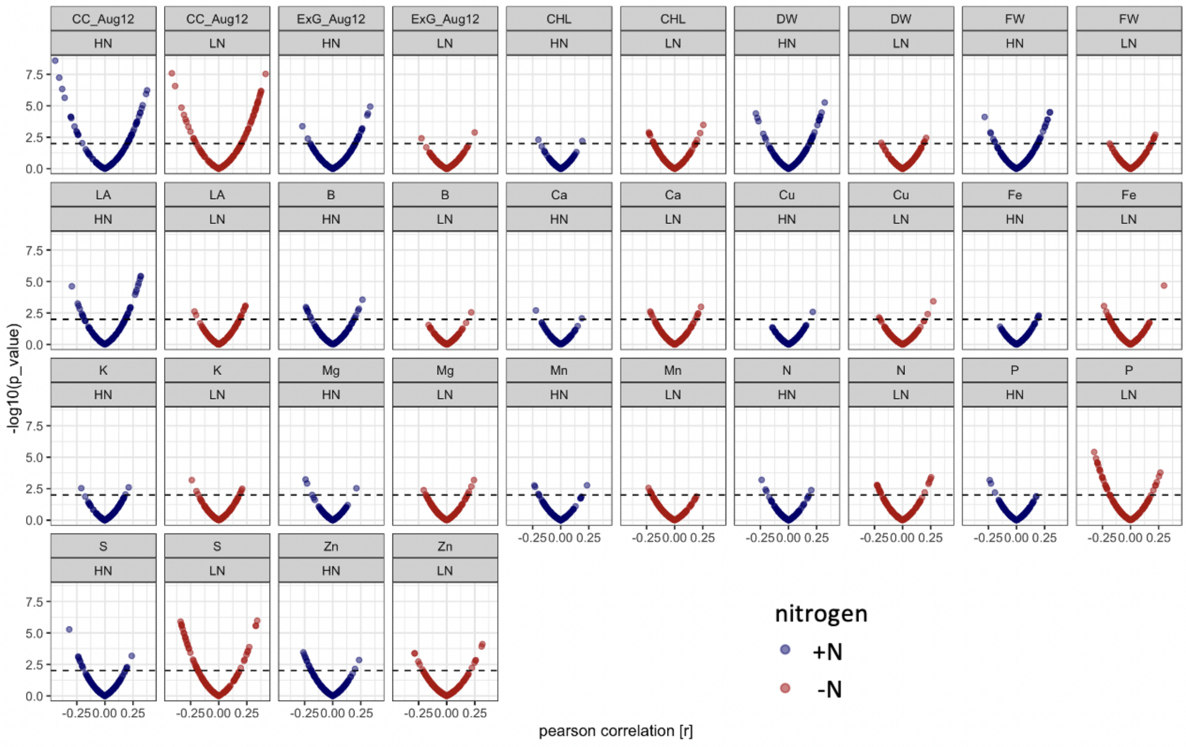


**Supplementary Figure 4: Correlation of microbe abundance with 17 agronomic and micronutrient traits under +N (blue) and -N (red) conditions**. Each dot represents one of 150 rhizobiome traits. X axis shows correlation with agronomic trait (r value), y axis shows significance, dashed line shows p=0.01 level of significance. CC_Aug12, EXG_Aug12: canopy coverage and excess green index measured on Aug. 12, 2019; CHL: chlorophyll content, DW: dry weight, FW: fresh weight, LA: leaf area.

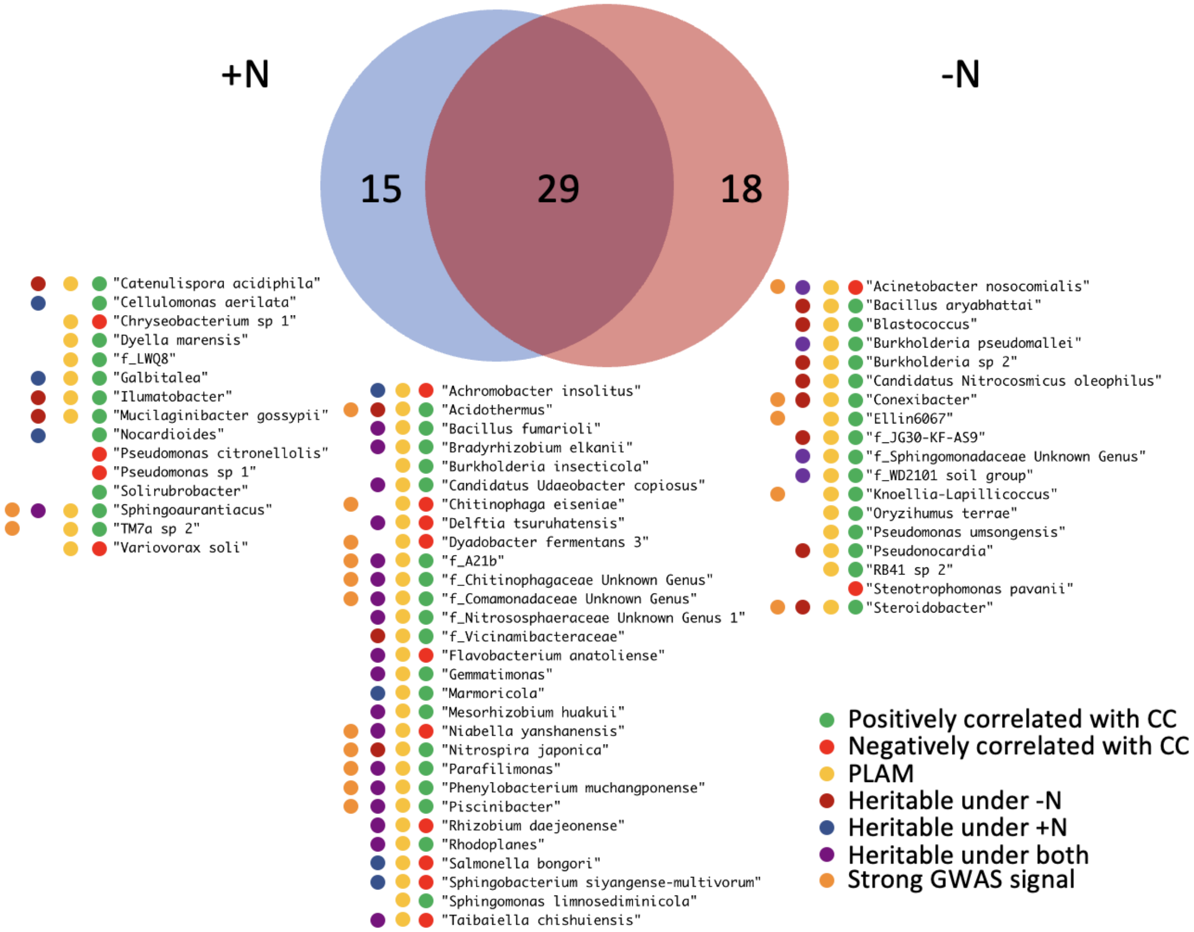


**Supplementary Figure 5: Microbial traits that correlate with canopy coverage.**
Venn diagram shows a total 62 microbial traits that correlate with canopy coverage either under +N, -N or both treatments. For the 62 listed rhizobiome traits, colored dots summarize various statistics that indicate association with the host plant.

A

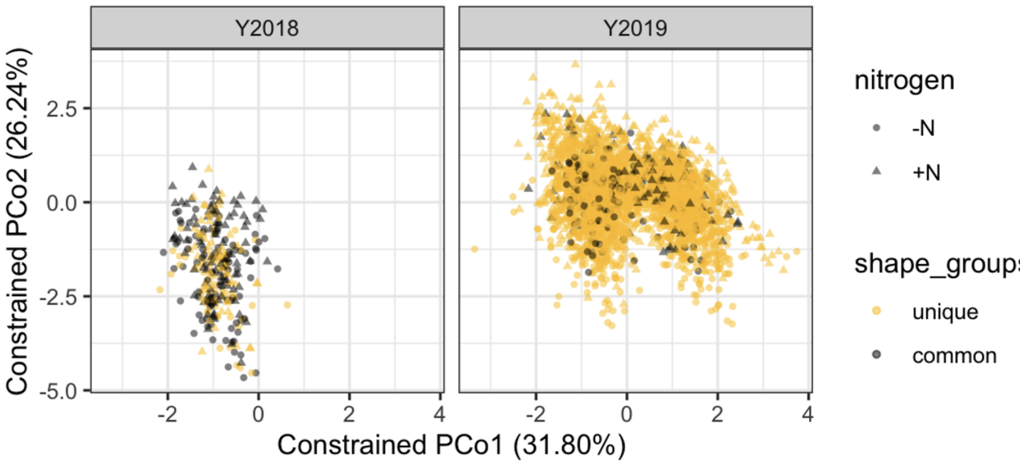


B

```
formula = dm ~ year + genotype + nitrogen + block + sp
```

| | Df | SumsOfSqs | MeanSqs | F.Model | R2 | Pr(>F) | |
|---|---|---|---|---|---|---|---|
| year | 1 | 3.354 | 3.3540 | 301.32 | 0.06311 | 0.001 | *** |
| genotype | 234 | 7.239 | 0.0309 | 2.78 | 0.13620 | 0.001 | *** |
| nitrogen | 1 | 1.783 | 1.7826 | 160.15 | 0.03354 | 0.001 | *** |
| block | 1 | 5.911 | 5.9105 | 531.01 | 0.11121 | 0.001 | *** |
| sp | 3 | 0.695 | 0.2317 | 20.81 | 0.01308 | 0.001 | *** |
| spb | 2 | 0.063 | 0.0314 | 2.82 | 0.00118 | 0.002 | ** |
| Residuals | 3064 | 34.105 | 0.0111 | | 0.64169 | | |
| Total | 3306 | 53.148 | | | 1.00000 | | |

**Supplementary Figure 6: PERMANOVA results.** It was calculated from the log(relative abundance) of 4,632 ASVs. Each dot represents a sample. Genotypes common to 2018 and 2019 panel are marked in grey.

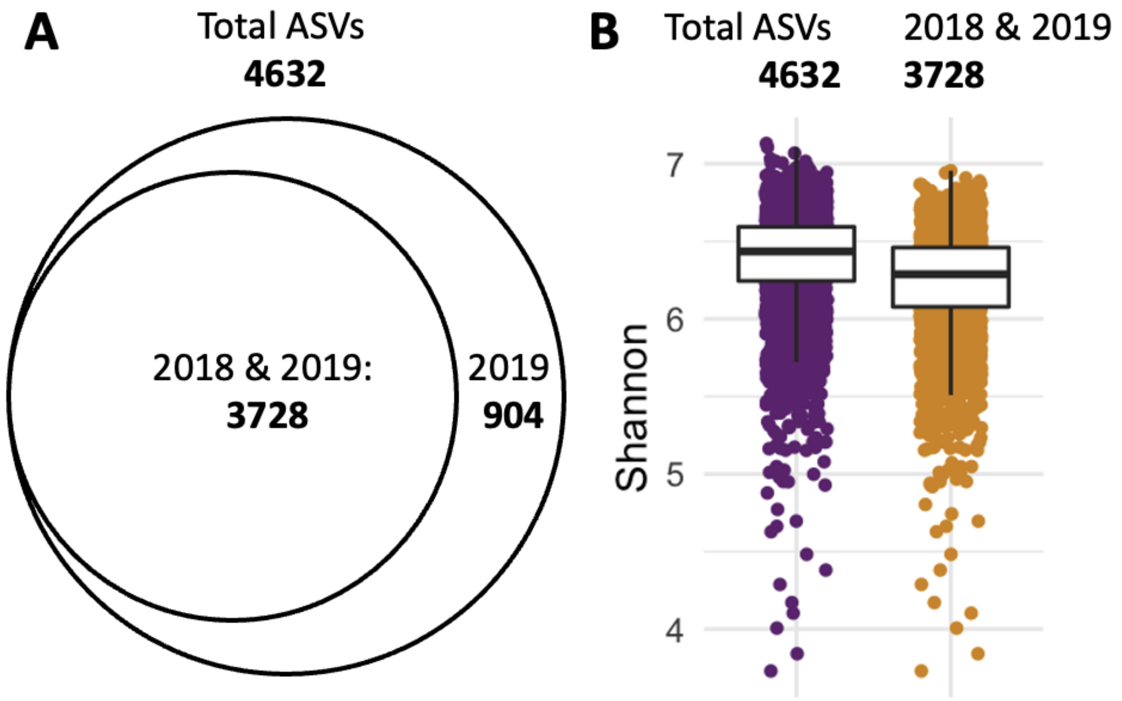


**Supplementary Figure 7**: **Retaining ASVs observed in both years reduces dataset complexity with minimal loss of diversity.** (A) Intermediate set of ASVs after prevalence filtering contains 4,632 ASVs, of which 904 were exclusively found in 2019. (B) Comparison of the Shannon diversity between the total set (4632 ASVs, purple) and the shared set (3728 ASVs, gold) reveals a 2.29% loss in diversity: Median(Shannon3728)/Median(Shannon4632) = 0.9771

### Supplementary Methods

##### Field and experimental Design

The experimental field was divided into 4 quadrants, which were separated and surrounded by a buffer of an industrial hybrid genotype (B73xMo17) (**Supplementary Figure 1**). The complete set of genotypes was planted in each quadrant where possible. Each quadrant was in turn divided into 4 split plots and a subset of the maize association panel was randomly assigned to each split plot based on the distributions of flowering time and plant height. Phenotypes were divided at the median value to create 4 flowering time / height categories: early/tall, late/tall, early/short, and late/short. Each split plot was further divided into 3 split plot blocks, and each split plot block was divided into 21 subplots in 3 ranges and 7 columns. Thus 252 subplots were available in each quadrant of the field. In each of 12 split plot blocks per quadrant, a t least one subplot was randomly selected and assigned the hybrid genotype (B73xMo17) to be used as a check to test for differences between geographical field locations. two check genotypes (B73xMo17 and B37xMo17) were used in 2018, and a single check genotype (B73xMo17) was used in 2019. Any subplots across the field that remained empty due to seed unavailability were filled with the check genotype as well.

In 2018, dry N fertilizer (urea) was applied to two diagonally opposed quadrants before planting at the rate of 140 kg/ha (+N treatment) while two quadrants were left unfertilized (-N treatment). In 2019, liquid N fertilizer (urea) was applied at the rate of 168 kg/ha. Both N treatments were thus represented in a northern block (NW and NE quadrants) and in a southern block (SW and SE quadrant). We assigned the blocks this way because of a 3 m increase in elevation from the north end of the field to the south end.

##### Rhizobiome sample preparation and sequencing

In 2018, rhizosphere samples were collected from 28 genotypes. These include, B73, the *roothairless3* mutant of B73 (Hochholdinger et al., 2008), two check hybrids (B73xMo17 and B37xMo17) and a subset of the Buckler-Goodman panel including 16 parent lines of the nested association mapping population (NAM) described by (McMullen et al., 2009). 8 weeks after planting, 2 subsamples per genotype were collected per quadrant and 12 subsamples for checks, where each subsample was taken from the combined root material of two adjacent plants. This resulted in a total of 26*4*2 +2*4*12 = 304 samples. In 2019, rhizosphere samples were collected in triplicates from all 1008 subplots within 3 days, 8 weeks after planting, when the majority of plants had reached the tasseling stage. One of the two rows in each subplot was randomly selected, and 3 individual randomly selected plants within the row (subsamples) were sacrificed for rhizosphere collection. As a small fraction of subplots had poor germination and/or no surviving plants on the day of sampling, the final number of rhizosphere samples collected was 3009. Rhizosphere samples were placed on ice immediately after collection and shipped to the lab to be processed on the same day.

To wash rhizosphere soil off the roots, tubes were filled up to the 40 ml mark with autoclaved PBS buffer (46 mM NaH2PO4, 60 mM Na2HPO4, 0.02% Silwet-77), and shaken horizontally at 8000 rpm for 30s. Rhizosphere suspension was filtered through a 100 µm nylon cell strainer (Celltreat Scientific Products, Pepperell, MA, USA) into a fresh 50 ml tube to capture root debris and large soil particles. Rhizosphere samples were frozen in suspension at -20°C until further processing. DNA was isolated from rhizosphere soil using the MagAttract PowerSoil DNA KF Kit (Qiagen, Hilden, Germany) and the KingFisher Flex Purification System (Thermo Fisher, Waltham, MA, USA) with minor modifications to the protocol: Rhizosphere samples that were kept in suspension were thawed on ice, pelleted soil was resuspended by inverting tubes, and 500 µl soil suspension was added to the 96-well sample plates. To avoid cross contamination of wells during pipetting, plates were sealed beforehand with parafilm and the cover was pierced with the pipette tip to transfer the rhizosphere suspension into the intended well. Two plates were prepared at a time and centrifuged for 10 min at 4000 x g to pellet soil. Supernatant was carefully removed with a multichannel pipette and 96-well plates with approximately 100-250 mg rhizosphere soil per well were frozen at -20°C until further processing. On the day of DNA isolation, the bead mill substrate was added to the frozen soil pellets, soil was thawed on ice and the remainder of the protocol was followed as per the manufacturer's instructions. We recommend this modified procedure for large numbers of samples as it is cleaner, faster, and better reproducible than scooping soil from pellets in sample tubes. Concentration of isolated DNA was measured fluorometrically with the QuantiFluor dsDNA System (Promega, Madison, WI, USA) as per the manufacturer's instructions. DNA isolation was repeated for any samples that failed to reach a concentration of at least 1 ng/µl.

A 350 bp stretch of 16S rDNA spanning the V4 region was amplified using V4_515F_Nextera (TCGTCGGCAGCGTCAGATGTGTATAAGAGACAGGTGCCAGCMGCCGCGGTAA) and V4_806R_Nextera (GTCTCGTGGGCTCGGAGATGTGTATAAGAGACAGGGACTACHVGGGTWTCTAAT) primers on several Illumina MiSeq runs. Oligonucleotide PCR blockers (PNA Bio INC, Thousand Oaks, CA, USA) targeting mitochondrial and chloroplast sequences were applied in the primary V4 amplification to reduce amplification of templates derived from the plant host. Up to 128 barcoded samples were pooled per sequencing run. In total, 304 samples in 2018 and 3009 samples in 2019 were sequenced on the same Illumina MiSeq machine.

**Raw read processing and construction of microbiome dataset**

Cluster computing resources at the UNL Holland Computing Center were used for computationally demanding steps. To construct the microbiome dataset, 350 bp raw sequencing reads were trimmed using *filterAndTrim()* at 240 bp (forward reads) and 200 bp (reverse reads), respectively. Amplicon sequence variants (ASVs) were inferred using *dada()* and forward and reverse reads were merged with *mergePairs()*. A sequence table was generated using *makeSequenceTable()* and chimaeras were removed using *removeBimeraDenovo()*. Taxonomy was assigned to ASVs with *assignTaxonomy()* using the SILVA database version 138 (Yilmaz et al., 2014) as a reference. SILVA was our taxonomy of choice because it is a relatively large 16S sequence database compared to alternative databases, it is regularly maintained and updated and it is widely used in ecological research, making our results comparable to other 16S studies. (Balvočiūtė and Huson, 2017). Taxonomic training data formatted for DADA2 (silva_nr99_v138_wSpecies_train_set.fa.gz) was obtained from https://zenodo.org/record/3986799#.X3zmypNKh24, as referenced by https://benjjneb.github.io/dada2/training.html on GitHub. 16S reads and sample data were prepared in an R Phyloseq object for further processing.

Raw ASV reads were subjected to a series of filters to produce a final ASV table with biologically relevant 16S sequences:

1) Removed chimaeric 16S reads using *removeBimeraDenovo*()
2) Removed sequences with <20 total observations
3) Removed sequences that did not map to either Bacteria or Archaea
4) Removed chloroplast sequences
5) Removed mitochondrial sequences
6) Removed ASVs that were not observed in at least 5% (166) of all samples
7) Removed ASVs that were not observed in both years 2018 and 2019
8) Removed 53 out of 160 genera and families that had fewer than 5 unique ASVs

4,632 common ASVs that were detected in at least 5% of the samples, representing 120,004,239 of the raw reads. Constrained ordination and PERMANOVA analyses of the 4,632 ASVs identified a strong effect of N treatment as well as other experimental factors on ASV abundance (**Supplementary Figure 6**). This observation is consistent with previous observations that environmental factors play an important role in determining the composition of the root associated microbiome diversity (Floc'h et al., 2020; Meier et al., 2021; Schlatter et al., 2020). Of the 4,632 common ASVs, 3,728 (or 80.5%) were highly abundant and observed in samples collected from both the 2018 and 2019 growing seasons. Removing ASVs that could not be repeatedly observed in multiple years reduced the complexity of the data set by 19.5% at the cost of a 2.3% loss in diversity (Shannon diversity reduced from 6.4 to 6.3, **Supplementary Figure 7**). Finally, removing taxa (genus or family) with less than 5 observed ASVs yielded a dataset of 3,626 ASVs, 3,313 samples, and 105,722,181 total ASV counts. This final core microbiome encompasses <1% of initial ASVs and ~50% of initial observations. The ASV table from step 8 was converted to relative abundances and values were transformed with the natural logarithm. A phylogenetic tree was constructed from the final set of 3626 ASVs using mafft v. 7.404 (Katoh and Standley, 2013) for multiple alignment and fasttree v. 2.1 (Price et al., 2010) and the phylogenetic tree was attached to the phyloseq object.

**Heritability estimation**

To calculate heritability ($h^2$), read counts from 3 subsamples were pooled for each subplot. Combined counts were then normalized by converting to relative abundance and subsequent natural log transformation, which yielded a subplot-level measure of microbial abundance, replicated in 2 blocks. The following linear mixed model was used with all random effects: **Y = genotype + block + error.** Y is the log-transformed relative abundance of each microbial group in each subplot-level sample, the blocks and subplots are as outlined in (**Supplementary Figure 1**). Heritability was tested for significance using a permutation test in which microbial abundance data for each trait was shuffled and heritability calculated anew 1000 times. p-values indicating heritability were calculated by tallying the number of permutation $h^2$ scores exceeding the observed $h^2$ and dividing by the number of permutations. Traits with a p-value < 0.05 were deemed “heritable” under either or both N treatments.

**Estimation of genetic architecture parameters**

SNPs in high linkage disequilibrium (LD) were pruned using the “indep-pairwise” command of with a LD threshold of r2 = 0.1. In the GCTB analysis, the BayesS model was used with the chain length of 410,000 and burnin 10,000. One example command used for the GCTB analysis is “gctb –bfile 282_GCTB_G --pheno gctb_blup_stdN_150_tax_groups.txt --mpheno 28 --out Results_HN/asv_000013 --bayes S --pi 0.05 --hsq 0.5 --S 0 --wind 0.1 --chain-length 410000 --burn-in 10000”.

**Genome-wide association study**

GWAS was performed using GEMMA 0.98 (Zhou and Stephens, 2012) with the following parameters: gemma-0.98 -bfile {snp_file} -k {kinship_matrix} -c {pca_file} -p {traits_file} -lmm 1 -n {trait_num} -outdir {outdir_path} -o T{trait_num} -miss 0.9 -r2 1 -hwe 0 -maf 0.01'). Blup values were summarized in a trait matrix (214 genotypes x 150 traits) for all 150 microbial traits and for all 214 maize genotypes for which high quality SNP data was available. To conserve disk space, SNP information was only retained in each ASV if a response at p_wald < $10^{-2}$ was observed. To identify genomic loci with high counts of significant SNPs, the genome was split into bins of 10 kbp, and the number of significant SNP signals at a threshold of p_wald < $10^{-5}$ was counted for each bin.
